# Combined impacts of invasive alien species and fire on ecosystems are complex, mostly negative, and understudied: a global review

**DOI:** 10.64898/2026.03.08.710355

**Authors:** Cristina G. Lima, Paulo Fernandes, Cândida G. Vale, João Gonçalves, João P. Honrado, Adrián Regos, Joana R. Vicente

## Abstract

Invasion-fire interactions can negatively impact ecosystems by driving biodiversity loss, altering ecological processes, modifying habitat structure, and compromising ecosystem functioning. Understanding how this interaction operates is essential to design effective management strategies that are successful in controlling both invasive alien species (IAS) and fire. Therefore, the present literature review aims to synthesize the current knowledge on invasion-fire interaction and its impacts on ecosystems, as well as identify knowledge gaps in the field. The review included 464 studies, from which information on context, species and fire characteristics, interaction outcomes, and research approaches was extracted. Fire generally promotes IAS, although studies on invasive animals are limited and no research has examined the effects of fire on fungi. Management through prescribed fire showed significantly better outcomes than wildfires in suppressing IAS, yet positive impacts still outnumbered the negative ones. In turn, IAS can change fire regimes causing regime shifts, but this direction of interaction is much less studied. Combined impacts of fire and IAS on ecosystems are predominantly negative, although interactions are complex and not always synergistic. Key knowledge gaps include geographic regions with known Invasion-fire interactions that remain underreported, a lack of broad-scale studies, limited management interventions, understudied taxa, and limited understanding of the combined effects on ecosystems. Remote sensing and laboratory experiments, which have been rarely used, could address some of these gaps.

## 1. Introduction

In ecology, a disturbance is a “relatively discrete event in time that disrupts ecosystem, community, or population structure and changes resources, substrate availability, or the physical environment” (White and Pickett, 1985). Disturbances regimes are fundamental components of ecosystems that shape their state and trajectory, yet they are changing rapidly due to global change (Turner, 2010). Drivers of change such as biological invasions can modify (wild)fire disturbance regimes (Brooks et al., 2004).

Invasive alien species (IAS) can change fire behaviour characteristics of an area by altering fuel characteristics (Mandle et al., 2011; D’Antonio and Vitousek, 1992). These changes, in turn, can benefit IAS by creating conditions that favour their establishment and spread (Zouhar et al., 2008), potentially resulting in a feedback loop known as the invasion–fire cycle (Wagner and Fraterrigo, 2015). The combined effects of fire and IAS can differ from their isolated (Menon et al., 2025), challenging ecosystem resilience to fire and resistance to invasions (Chambers et al., 2014). This interaction can drive biodiversity loss, alter ecological processes, modify habitat structure, and compromise ecosystem functioning (Bowman et al., 2014; Le Maitre et al., 2014; Fisher et al., 2009; Rossiter-Rachor et al., 2008).

The growing concern over IAS and fire is reflected in policy and legislation, though these threats are usually addressed separately. Key measures for IAS management include, for example, the European Union’s Regulation 1143/2014 and the United States’ Executive Order 13112 (1999), while fire management has been the focus of instruments such as the European Union’s Wildfire Prevention Action Plan (2022) and the United States’ Wildfire Prevention Act of 2025. However, policy and legislation that explicitly integrates invasion-fire interactions remains scarce. An exception is the United States’ Secretary of the Interior Order No. 3336 (2015), which seeks to prevent and suppress rangeland fires while restoring sagebrush ecosystems degraded by invasive grasses.

Understanding the outcomes of invasion-fire interactions is essential for designing effective management strategies that are successful in controlling both IAS and fire risk (Hart-Fredeluces et al., 2025; Menon et al., 2025). Focusing on only one of these stressors has, in some cases, led to poor management outcomes. For example, McGlone et al. (2009) found that thinning and prescribed burning increased native understory plant diversity and cover but also facilitated invasion by the nonnative grass *Bromus tectorum* L. in ponderosa pine forests at Grand Canyon-Parashant National Monument (Arizona, USA).

In recent years, research on invasion–fire interactions has also increased substantially, with several studies and reviews published. Some recent reviews synthesize how invader growth forms affect the mechanisms by which and habitats in which invasion–fire cycles occur (Tomat-Kelly and Flory, 2022) and how fire can be used to control invasive woody species globally (Brancatelli et al., 2024). However, to our knowledge, no review has specifically focused on the ecosystem impacts resulting from these interactions. Addressing this gap is critical to inform management strategies in areas where IAS and fire interact, as well as in areas with high fire proneness and invasibility where such interactions may occur in the future. Therefore, the present review aims to synthesize the current knowledge on invasion-fire interaction and its impacts on ecosystems, identify knowledge gaps, and discuss possible future steps. The present study is organized into four sections addressing: (1) invasion-fire feedbacks, examining how IAS and fire affect each other; (2) impacts of IAS and fire on ecosystems, identifying the types of impacts caused and the affected habitats; (3) research approaches used to analyse the impacts of IAS and fire on ecosystems, focusing on the methodology used, aims of the studies, and result types present; and (4) an overview of main findings, key knowledge gaps, and future directions in the field.

## 2. Methods

The literature review followed a two-step framework based on standard guidelines (O’Dea et al., 2021), consisting of a literature search and review. The review was registered and carried out using the online tool CADIMA (Kohl et al., 2018), except for duplicate record removal which was performed in Excel.

The literature search was supported by a set of keywords related to “biological invasions”, “fire regimes”, and “interaction” (available in Appendix 1). Their selection considered keywords from literature related to the topic and a participatory approach with a team of researchers specialized in IAS and fire ecology. The full string of keywords can be found in Appendix 1.

In February 2025, searches were conducted using Scopus and ISI Web of Knowledge engines, resulting in a total of 1345 studies from 1984 (the date of the first study found) to 2024. Studies available online in 2024 and assigned to printed issues in 2025 were also included. To evaluate the reliability of the search, the dataset was compared to the top 50 records retrieved from Google Scholar using a simple string of keywords related to the topic (“invasive” AND “fire” AND “effect”). Duplicated studies were automatically removed by identifying matching digital object identifiers (DOIs). For studies without a DOI, duplicates were identified and removed manually by comparing titles.

Following the initial search, the screening process was performed to exclude irrelevant studies by applying inclusion/exclusion criteria. To be included, each study had to meet the following criteria: (i) address the interaction between invasive alien species and fire, (ii) be published in peer-reviewed literature, and (iii) present original research. These criteria were checked by evaluating the title, abstract, and keywords of each study, followed by a review of the full text. The final dataset included 464 studies in total, 11 of which were identified through the Google Scholar search. Appendix 2 shows all the studies gathered in the literature search and the final dataset of studies.

Once the final dataset was established, each study was thoroughly reviewed and classified according to a predefined set of categories (Figure 1), organized into four main groups to capture different aspects of the studies: (1) study context, (2) IAS and fire traits addressed, (3) the direction of invasion–fire interaction and its outcomes, and (4) the research approach of the study.

**Figure 1.**
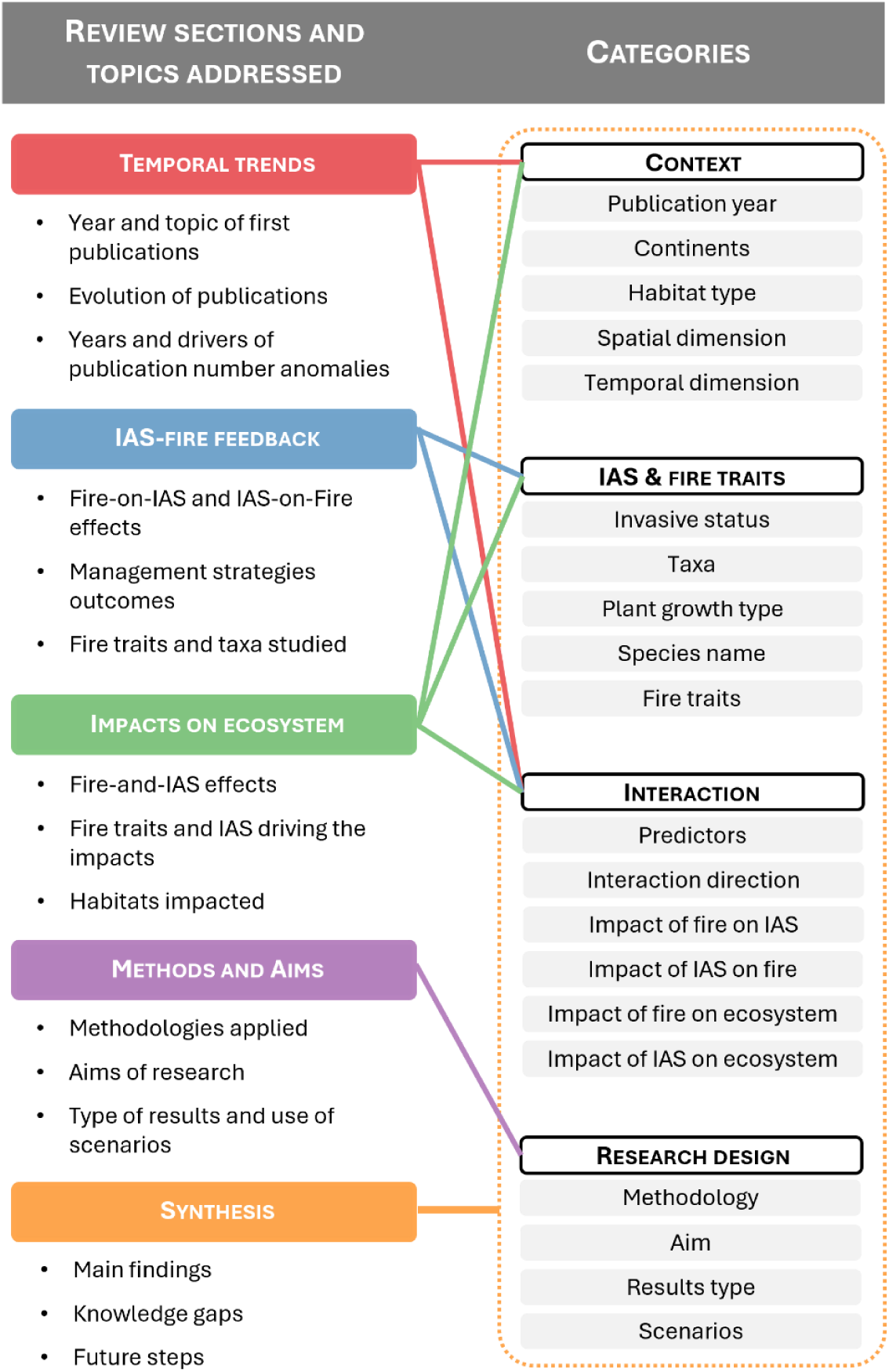
Conceptual diagram of the review structure and categories used for the data extraction. The left column shows the review sections and the main topics addressed in each. The right column lists the categories extracted from the reviewed studies, grouped into four category groups: Context, IAS & fire characteristics, Interaction, and Research approach. Coloured lines connect each review section to the category groups from which information was retrieved.

In the Context group, year of publication, continent, habitat type, and spatial and temporal scales were extracted. In the IAS and fire traits group, invasive status, taxonomic group, plant growth form (when applicable), scientific names of IAS, and fire regime attributes studied were extracted. In the Interaction group, predictors studied, direction of the invasion–fire interaction, effects of fire on IAS, effects of IAS on fire, and combined effects on ecosystems were extracted (effects per direction addressed only when applicable). Studies with multiple predictors and response variables were included, but our focus was on the direct interaction between fire and IAS and their combined ecological impacts. Finally, in the Research approach group, the methodology used, target aims, use of scenarios, and type of results presented were extracted. A full description of all categories is provided in Appendix 3, along with definitions for the categories fire regime attributes, model types, methodologies, and result types.

To facilitate clarity throughout the review, key terms used in this study are defined here. Studies about invasion-fire interactions were organized into three directions of interaction: *Fire-on-IAS*, referring to the effects of fire on IAS; *IAS-on-Fire*, referring to the effects of IAS on fire; and *Fire-and-IAS*, referring to the combined effects of fire and IAS on ecosystems. These terms are used to condense lengthy expressions and enhance readability. Throughout this review, the terms *study* and *record* are used. *Study* refers to a scientific article included in the dataset and a *record* refers to each unit of information extracted from a study within the predefined categories. In other words, when a study provides more than one entry in a given category, it generates multiple records.

To explore patterns in the dataset, univariate and statistical analyses were performed. Univariate analyses included line plots, bar plots, and multi-layer pie donut plots. Statistical analyses included Chi-square and Fisher’s exact tests. All analyses were conducted in Python using the libraries Matplotlib and SciPy.

## 3. Temporal evolution of invasion-fire research

The first study about invasion-fire interactions was published in 1987 and focused on Fire-on-IAS, examining the impacts of clearfelling and slash-burning on vascular plants in two Eucalypt forests in Tasmania (Dickinson and Kirkpatrick, 1987). The first study addressing Fire-and-IAS was published two years later. It explored how managing *Hakea sericea* Schrad. & J. C. Wendl through cutting and burning after seedling emergence negatively affected the fynbos ecosystem of South Africa (Breytenbach, 1989). The first study on IAS-on-Fire was published in 1998, showing how grass invasion in woodlands altered modelled fire behaviour (Freifelder et al., 1998).

After these initial studies, publications on the topic appeared sporadically and in small numbers until 2005, when the first notable peak occurred, followed by a steady stream of research (Figure 2). This 2005 peak is largely driven by an increase in Fire-on-IAS studies. It may reflect a growing awareness among researchers, likely influenced by the publication of several impactful reviews in preceding years. More than half of the records cite at least one of the following works: Brooks et al., 2004; Levine et al., 2003; D’Antonio and Meyerson, 2002; D’Antonio et al., 1999; D’Antonio and Vitousek, 1992.

**Figure 2.**
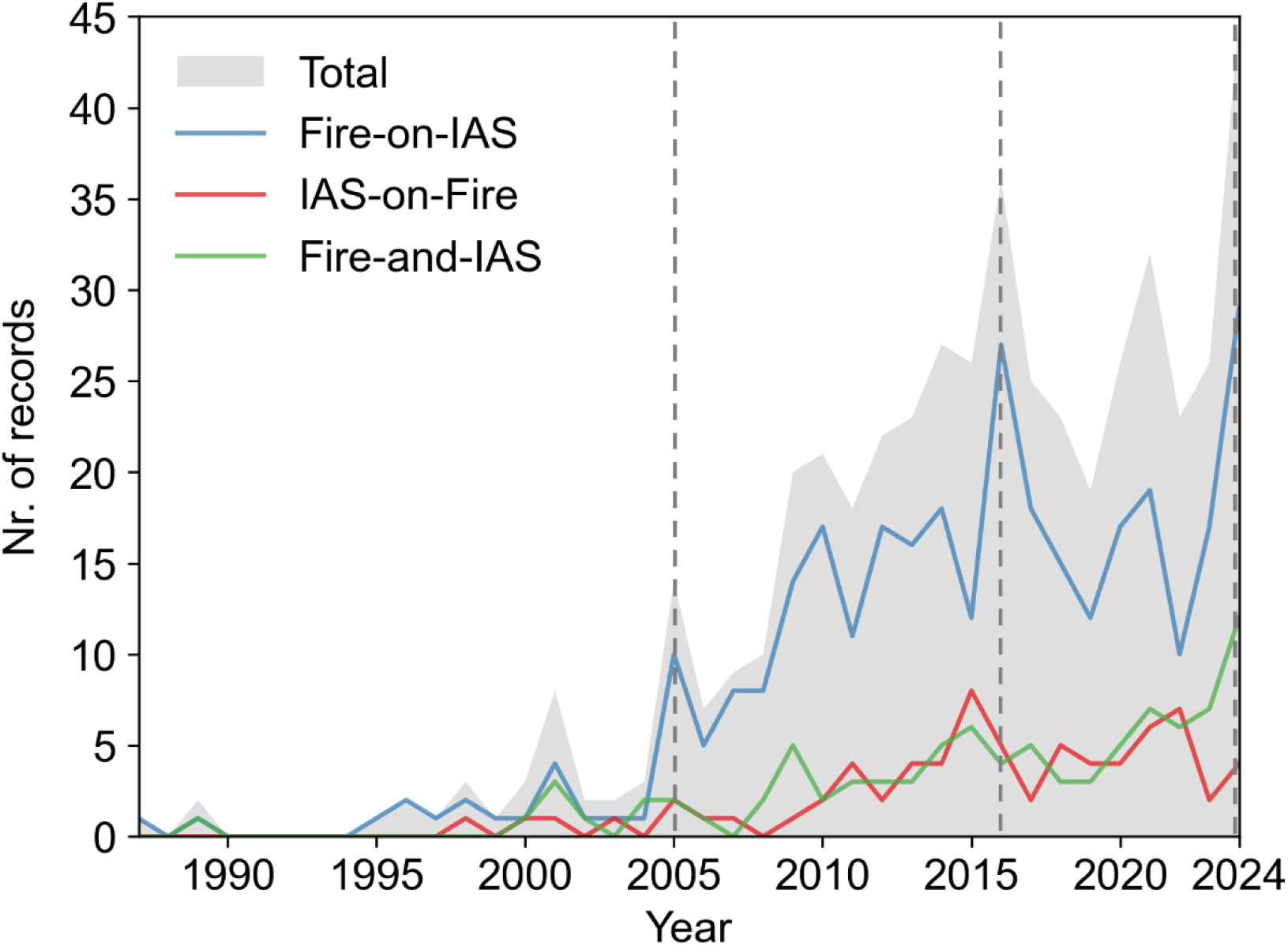
Annual number of published records addressing the interaction between fire and invasive alien species (IAS) from 1984 to 2024. The total number of records is shown as a filled grey area. Records focusing on fire effects on IAS (Fire-on-IAS) are shown in blue, records examining IAS effects on fire (IAS-on-Fire) in red, and records addressing the combined effects of fire and IAS on ecosystems (Fire-and-IAS) in green.

After 2005, IAS-on-Fire and Fire-and-IAS had a slow and slight increase in the number of publications per year. Fire-on-IAS displayed a much more marked increase, with a distinct peak in 2016 and reaching a record high in 2024 (Figure 2).

In 2016, all the studies reviewed framed their research around the impacts and management of IAS, with most focusing their research questions on testing potential management strategies. This trend reflects a broader rise in concern about IAS, evidenced by an increase in both scientific publications and legislative action, as well as a growing emphasis on biodiversity conservation. For instance, in 2015, the United States’ Secretary of the Interior issued Order No. 3336, which aimed to prevent and suppress rangeland fires and restore sagebrush landscapes. One of its goals was to reduce the likelihood, size, and severity of rangeland fires by addressing the spread of IAS and mitigating threats from IAS. This aligns with the scientific publications, as the majority are from the American continent and many discuss the protection of sagebrush ecosystems from IAS, like *B. tectorum* (e.g., Gray and Dickson, 2016; Porensky and Blumenthal, 2016; Reed-Dustin et al., 2016).

The year 2024 represents the highest publication output recorded in the field, with the focus on management continuing but showing notable shifts. Firstly, conservation concerns, together with the management of IAS and fire, have gained significant political attention, leading to a surge of restoration projects, species protection initiatives, and new strategic plans. This seems to be orienting studies on this topic into conservation aims. Concern over the impacts of IAS and the need for management continues to grow, shaping the focus of many of the recent studies on the topic (e.g., Alba et al., 2024; Giles et al., 2024; Ngwenya et al., 2024; Urza et al., 2024). In addition, the impacts of changing fire regimes and the increasing occurrence of ‘megafires’ are causing alarm and being addressed by policymakers (e.g., UNEP, 2022; BLM, 2020), a trend highlighted in several studies (e.g., Davies et al., 2024; Dickson-Hoyle et al., 2024; Germino et al., 2024; Kraaij et al., 2024). Finally, references to restoration and conservation plans are also at the forefront of framing recent studies (e.g., DCCEEW, 2023; UNEP and FAO, 2020; USDA Forest Service, 2018). Secondly, an increasing number of studies discuss the limitations of existing management strategies, reporting ineffectiveness or even unintended negative impacts (e.g., Price et al., 2025; Davies et al., 2024; Ngwenya et al., 2024). For example, since *Vulpes vulpes* L. tend to hunt native mammals more effectively after fire due to the loss of understory cover, Menon et al. (2025) investigated whether fox baiting programs could mitigate these effects of prescribed fire. The study found that fox occupancy increased after fire across all sites, but the increase was smaller in areas where baiting was applied. However, in these baited areas, *Felis catus* L. occupancy doubled. Therefore, this strategy did not improve the short-term resilience of native mammals to fire. In another study, Vorup and Pagter (2024) concluded that prescribed fire, used to manage Danish heathlands, was promoting the germination and seedling emergence of the exotic *Cytisus scoparius* L.

It is also noteworthy that this research topic is expanding, with increasing diversity in the methodologies applied (e.g., remote sensing), the taxonomic groups investigated (e.g., animals), and the geographic regions explored (e.g., Europe), many of which had previously received little attention. These elements have gained momentum in recent years, reaching their highest levels in 2024 (see Appendix 4).

## 4. Invasion-fire feedback

On Fire-on-IAS records, herbaceous plants are the most examined taxonomic group, followed by trees and shrubs. Only 16 records (out of 395) have analysed invasive animals, and no research has addressed the effects of fire on invasive fungi (Figure 3A).

**Figure 3.**
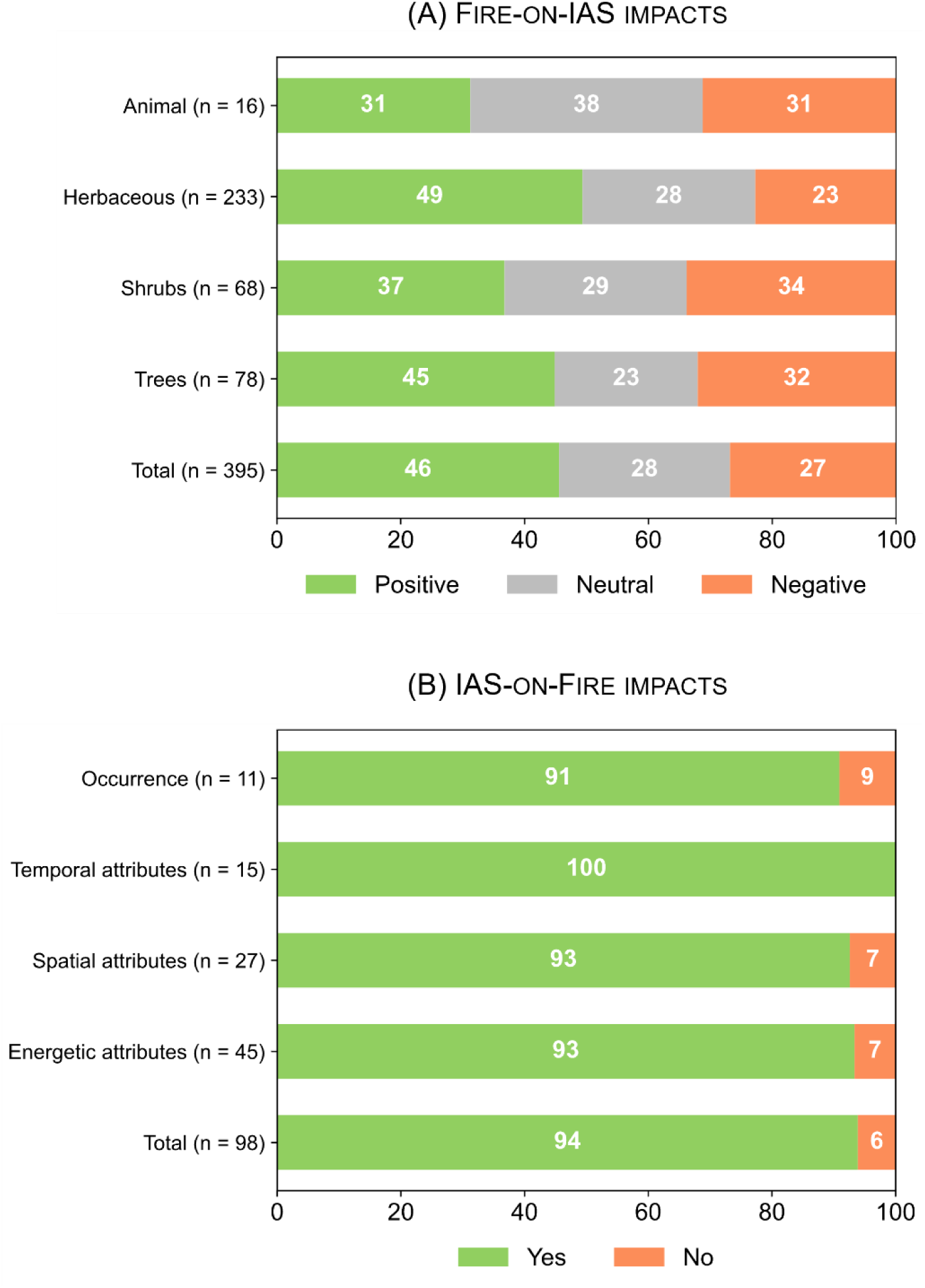
Stacked bar plots showing the interaction between fire and invasive alien species (IAS), divided in two panels. (A) Fire-on-IAS impacts, grouped by taxonomic group. Positive impacts shown in green, Neutral in grey, and Negative in red. (B) IAS-on-Fire impacts, grouped by fire attributes studied. Fire behaviour or regime changes are shown in green, and no changes in red.

Fire can promote IAS across all taxa. The number of records reporting positive impacts exceeds those reporting negative impacts for all IAS taxa, except for animals, where positive and negative reports are equally frequent (Figure 3A). The positive impacts on invasive plants can be attributed to the window of opportunity that fire creates for invasion, influencing the three primary factors that determine their establishment: resource availability, species traits, and propagule pressure (Brooks, 2008). Fire increases resource availability by reducing occupancy and resource use by the pre-fire vegetation (e.g., causing mortality) or by altering the form and availability of nutrients, light, and water (Cavallero and Raffaele, 2010; Brooks, 2008). Fire can also change the balance between native species and IAS, since species can be either fire-sensitive or fire-adapted (e.g., Trouvé et al., 2020; Bond and Keeley, 2005; Brooks et al., 2004). Depending on the post-fire response of the native and IAS present in the ecosystem, fire can change this balance and promote IAS. Finally, fire effects on propagule pressure depend on species-specific traits. It can kill seeds in heat-sensitive species or promote germination by breaking seed dormancy (Suárez-Ronay et al., 2024). The mechanisms through which fire benefits invasive animals are not as well studied. From the few studies focused on invasive animals that report positive impacts, benefits occurred through two mechanisms. First, fire increases resource availability (e.g., open space), which IAS better adapted to post-fire conditions exploit (Malumbres-Olarte et al., 2014). Second, and more commonly reported, the post-fire reduction in vegetation cover increases prey vulnerability and facilitates predation by IAS (Trewella et al., 2023; Bliege Bird et al., 2018; Hradsky et al., 2017).

Studies about IAS-on-Fire are much less frequent than Fire-on-IAS (Figure 3). Most IAS-on-Fire records report changes in fire regime, across all fire characteristics (Figure 3B).

Fire regimes are shaped by the interactions among topographic patterns, climatic conditions, ignition sources, and fuel characteristics (Rego et al., 2021). Vegetation fuels can change in days or weeks due to land cover conversion to agriculture or other disturbances, or years to decades due to plant invasion, which can displace native vegetation and change community or landscape-level flammability (Brooks, 2008).

Alien plants can directly change flammability, through their fuel properties, or indirectly by altering the structure of native fuels (Brooks, 2008). For example, Cubino et al. (2018) examined the influence of morphological and flammability traits of native and exotic plant species on community-level flammability in tussock grasslands over a 25-year period. They concluded that invasion by exotic forbs reduced overall flammability, potentially leading to lower-intensity, patchier fires and long-term changes in plant community composition. Similarly, McGranahan et al. (2013) analysed how the invasion of a high-moisture cool-season grass, *Schedonorus phoenix* (Scop.) Holub, affects fire spread in tallgrass prairies in Iowa (USA). The authors concluded that this IAS reduces fire spread by increasing live fuel moisture and fuel discontinuity, showing that invasive grasses can dampen rather than intensify fire regimes. Regarding changes in fuel structure, Paritsis et al. (2018) evaluated changes in the structure and potential fire behaviour across plots of native steppe, pine plantations, pine invasions, and removed invasions in Patagonia. They concluded that plantations and invaded areas changed fine fuel loads and their vertical and horizontal continuity, increasing fire intensity.

Invasive alien species can quantitatively affect the fire regime reflecting changes in the magnitude of pre-existing fuel (typically less impactful) or qualitatively reflecting a change in fuel type (Brooks, 2008). For instance, in the Cerrado biome of Brazil, invasion by African grasses accumulate more biomass than native herbaceous species, making them the primary combustible plants and increasing the severity of fires (Rossi et al., 2014). On the other hand, in the northern Great Basin, invasion of sagebrush ecosystems by exotic annual grasses such as *B. tectorum* and *Taeniatherum caput-medusa* Nevski changes the fire regime qualitatively (Fernández-Guisuraga et al., 2023).

In Fire-on-IAS, the impacts of wildfires and management on IAS through prescribed burning were compared using a chi-square test. The two fire types affected IAS differently (χ^2^ = 7.73; p-value < 0.05), with prescribed burning records reporting fewer IAS promotion and more neutral or negative impacts (Figure 4). Nonetheless, positive impacts on IAS remained the most common outcome for prescribed burning, occurring in 37% of cases. For example, Sabo et al. (2009) addressed the effects of thinning and prescribed burning on understory plants in ponderosa pine forests of northern Arizona. Thinned and burned stands contained 7–11% non-native grasses, compared to none in unmanaged stands. In another study, Pavlovic et al. (2016) concluded that invasive *Celastrus orbiculatus* Thunb. is highly responsive to fire disturbance and that fire can facilitate its spread in the oak savannas of Indiana Dunes National Lakeshore by stimulating vigorous resprouting and root-suckering.

**Figure 4.**
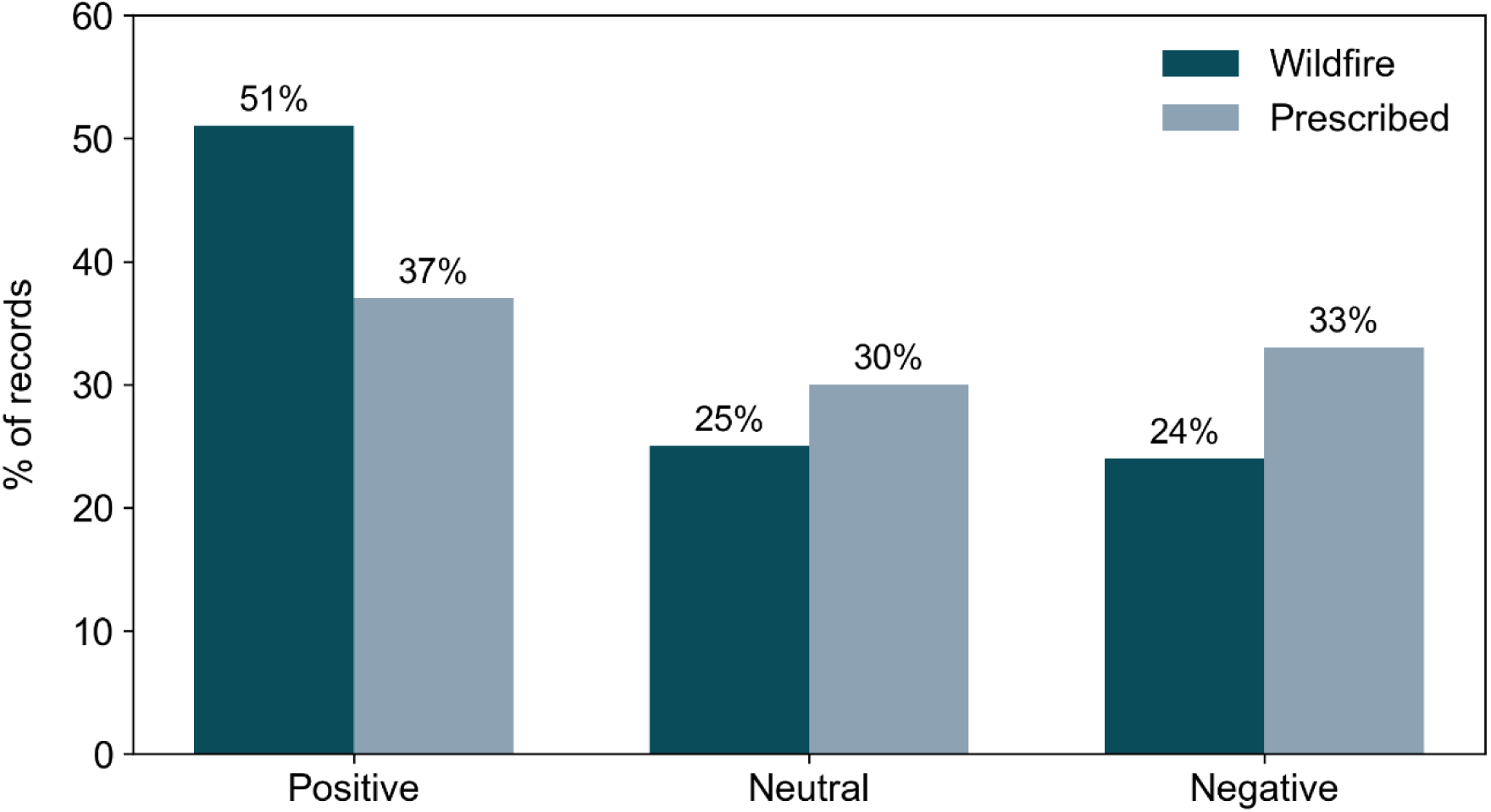
Bar plots showing the proportion of Positive, Neutral, and Negative impacts of wildfires and prescribed fires on invasive alien species.

The limited number of studies examining the effects of IAS management on fire regimes hampered a comparable analysis for IAS-on-fire.

A subset of 51 records within the dataset examined the interaction between fire and alien species rather than focusing specifically on IAS. These records were excluded from the main analyses, as the review specifically addresses Invasion–fire interactions. Nevertheless, this subset—dominated by records on the effects of fire on alien species (n = 41) — shows that fire can facilitate the transition of alien species into problematic invaders (e.g., Keeley and Brennan, 2012; Keeley et al., 2011). Among these records, 46% reported positive impacts of fire, 29% neutral impacts, and 25% negative impacts. Fire benefits alien species through similar mechanisms as IAS. Some records address fire benefits to alien species through changes in ecosystem conditions and resource availability (e.g., Crawford et al., 2001), whereas others show advantages linked to species traits (e.g., Clarke and French, 2005). Management interventions can also inadvertently promote alien species with, for example, mastication treatments used to reduce fire hazard increasing alien frequency and cover by altering canopy and ground conditions (Coop et al., 2017). These findings highlight that fire can facilitate the transition of alien species into problematic invaders, underscoring the need for management strategies to consider both established invaders and alien species.

## 5. Combined impacts of invasion-fire interactions on ecosystems

Invasion-fire interactions most often have negative impacts on ecosystems (Figure 5). There is no significant difference between the impacts of fire and IAS on the ecosystems (χ² = 4.55; p-value = 0.1). Furthermore, neither fire nor IAS impacts on ecosystems differed significantly among habitats (fire: odds ratio = 1.46 × 10⁻⁸, p-value = 0.36; IAS: odds ratio = 6.15 × 10⁻⁷, p-value = 0.47). This indicates that the impacts of these drivers are not habitat-specific but rather broadly distributed across the studied systems. However, published literature suggests that habitats can differ in their resistance to invasion (Chytrý et al., 2008) and in their resilience to fire (Mitchell et al., 2009). A possible explanation is that research has focused on habitats where invasion-fire interactions impacts are more pronounced, since these require particular attention for conservation and, as discussed in the Temporal trends section, recent priorities in this field are guided by conservation and restoration plans. The published work may thus reflect a subset of ecosystems that are vulnerable, leading to an apparent uniformity of impact across habitats.

**Figure 5.**
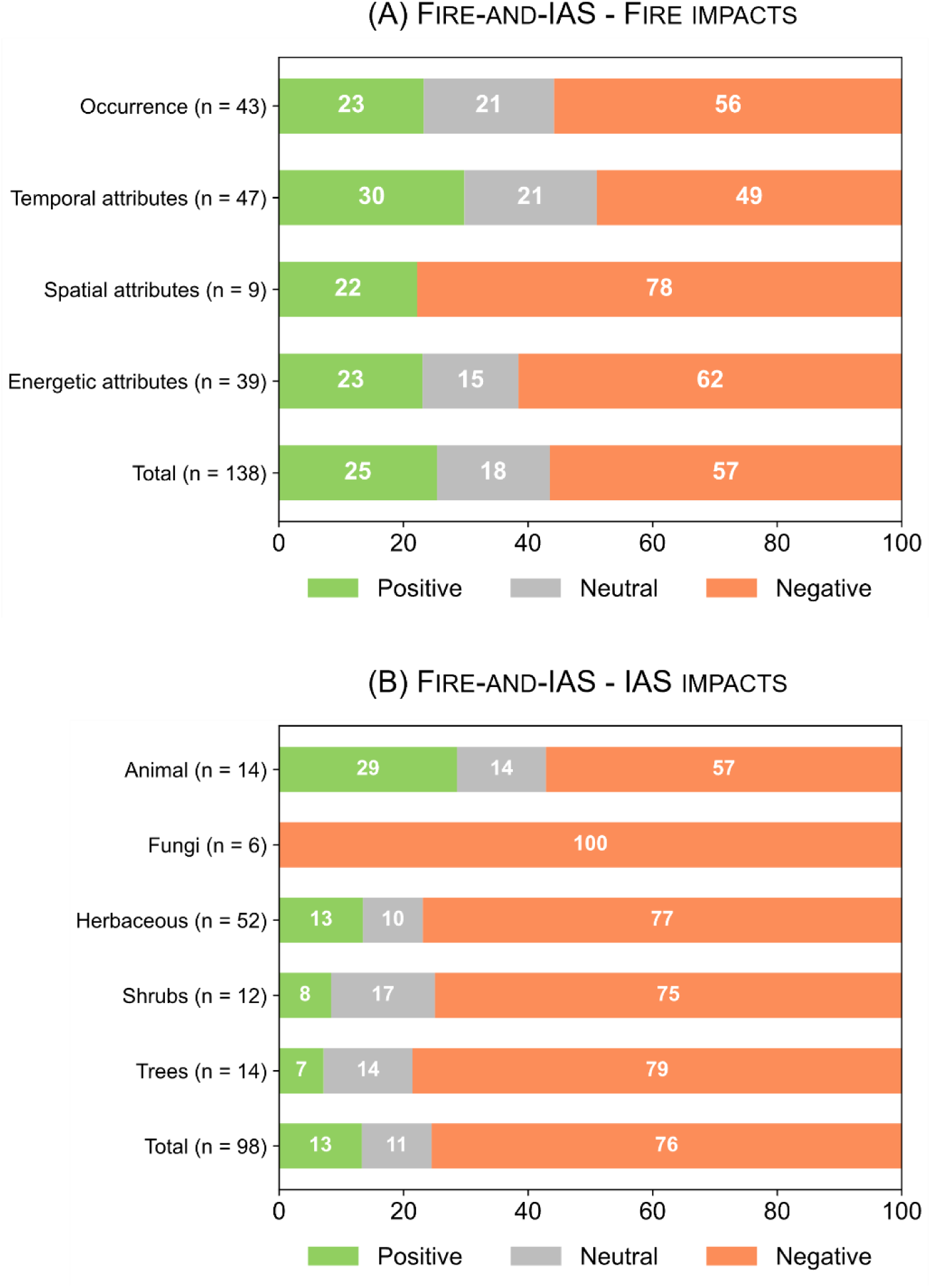
Stacked bar plots showing the combined impacts of fire and invasive alien species (IAS) on ecosystems (Fire-and-IAS), divided into two panels. (A) Impacts of fire on ecosystems, grouped by fire attributes. (B) Impacts of IAS on ecosystems, grouped by taxonomic group. Positive Impacts are shown in green, Neutral in grey, and Negative in red.

The most frequently studied fire traits are temporal attributes (n = 47), while spatial attributes are addressed in far fewer records (n = 9). Across all traits, the reported impacts are predominantly negative, although the extent of these impacts varies among attributes (Figure 5A). Records analysing spatial attributes report the highest proportion of negative impacts (78%), whereas those focusing on temporal attributes report the lowest (49%). These negative impacts are often linked to direct injury or mortality of native species, as well as shifts in community composition, such as reductions in richness and native cover (e.g., Macdougall et al., 2013; Fisher et al., 2009; Holmes, 2001). Several records also focus on changes in soil properties (e.g., Rossiter-Rachor et al., 2008; Mack et al., 2001; Breytenbach, 1989).

Nonetheless, a considerable number of records also report positive impacts of fire on ecosystems. These impacts can occur in diverse forms, including reducing the spread of disease (He et al., 2021), promoting community composition dynamics (Watson et al., 2021), and controlling IAS (Kaul et al., 2023). Nonetheless, the interaction between fire and IAS can alter the effects of fire on ecosystems, producing effects that differ from those observed in areas without invasion (Steidl and Litt, 2009). Therefore, management strategies must carefully account for the presence of IAS, the characteristics of the invaded community, and the conservation targets at stake, to design treatments that align with the targeted aims (Pyke et al., 2010).

The most frequently examined IAS taxonomic group in Fire-and-IAS is herbaceous plants, followed by animals, trees, and shrubs (Figure 5B). All plant groups show similarly high levels of negative impacts, whereas animals exhibit fewer negative impacts. The most frequently reported IAS with negative impacts is *B. tectorum*, followed by *Cenchrus ciliaris* L. and *V. vulpes* (see complete list in Appendix 5). Fungal IAS are analysed in only six records, and all report negative impacts (e.g., He et al., 2021; Loehman et al., 2017; Van Mantgem et al., 2004). These records focus on two fungi species, *Phytophthora ramorum* Werres, De Cock & Man and *Cronartium ribicola* J. C. Fisch.

Studies reporting negative impacts of IAS on ecosystems most often focus on their role in altering fire regimes (e.g., Franzese et al., 2020; Roldan-Nicolau et al., 2020; Rossiter-Rachor et al., 2008), which in turn leads to higher mortality and reduced recruitment of native individuals (e.g., Rahlao et al., 2009; Cilliers et al., 2004; Van Mantgem et al., 2004), along with decreases in the abundance and cover of native species (e.g., Geyle et al., 2025; Brehme et al., 2023; McDonald and McPherson, 2011). Several records also document shifts in vegetation composition (e.g., Dezotti et al., 2024; Le Maitre et al., 2014; Fisher et al., 2009), limitations to the post-fire recovery of native species (e.g., Salemme and Fraterrigo, 2021; Ramirez et al., 2012; Ainsworth and Kauffman, 2010), and reductions in species diversity (e.g., Tortorelli et al., 2020; Goerzen et al., 2019; Macdougall et al., 2013).

Most of the positive impacts of IAS on ecosystems are reported in records focusing on IAS management (e.g., Watchorn et al., 2024; Thomson et al., 2021; Bleicher and Dickman, 2020), although a few records also document the direct positive impacts of IAS. For example, De Marco et al. (2023) concluded that *Robinia pseudoacacia* L. improved soil content and availability of the studied mineral elements, as well as stimulated microbial growth. In another study, Steidl and Litt (2009) analysed the effects of fire and IAS on three native species. They found that the invader *Eragrostis lehmanniana* Nees had mixed effects: one native species was unaffected, one declined in abundance, and one increased.

## 6. Methods, aims, and result types in studies about invasion-fire impacts on ecosystems

Regarding methodologies used (Figure 6 inner rings), experimental studies are the most common (38%), followed by modelling (23%) and observational studies (19%), highlighting the strong reliance on field data (57%). Indeed, fieldwork is essential to generate empirical data on complex, context-dependent interactions such as invasion-fire interaction (Soga and Gaston, 2025). On the other hand, laboratory experiments and remote sensing are used less frequently, representing only 10% and 9% of the studies, respectively. Although less frequently used, these approaches have particular advantages. Laboratory experiments provide detailed insights into processes such as germination and plant growth (e.g., Mero et al., 2023; Calvo et al., 2013; Lopez-Arcos et al., 2012), while remote sensing enables analyses across large spatial and temporal scales (e.g., Rees et al., 2024; Visser et al., 2016; Setterfield et al., 2013).

**Figure 6.**
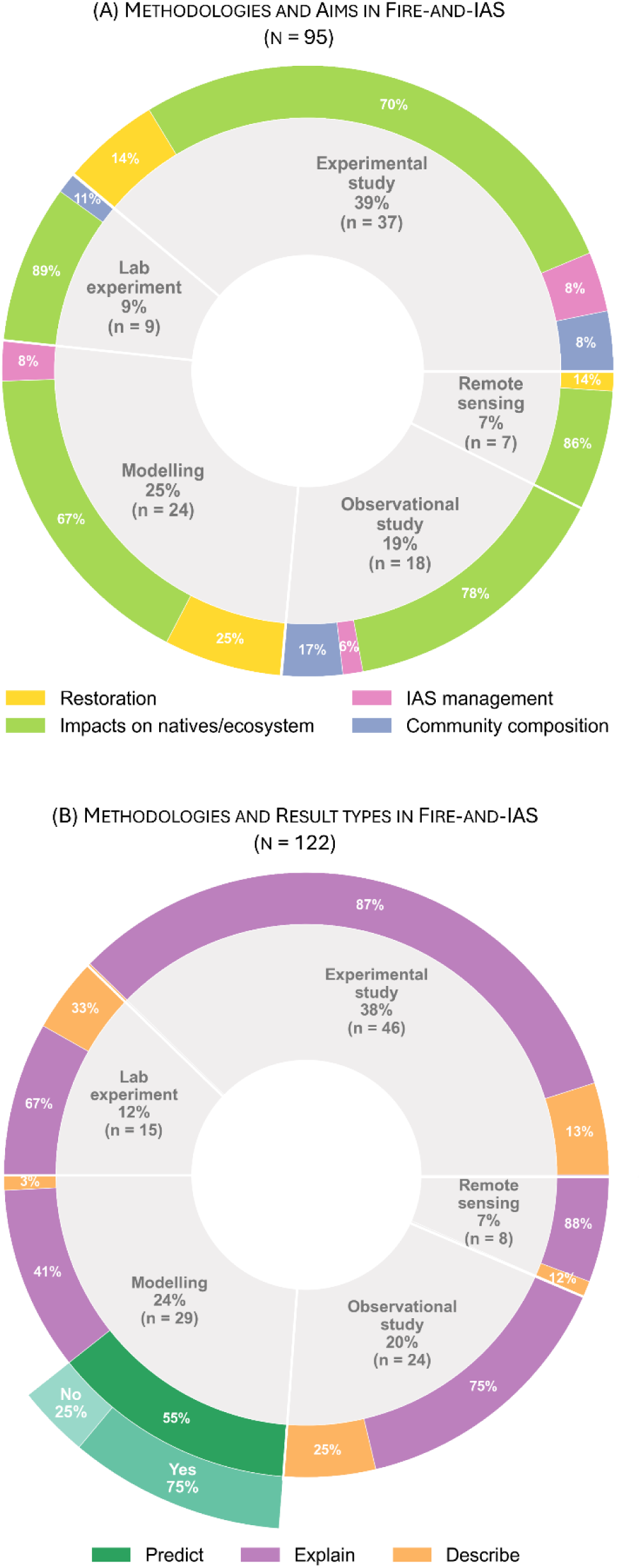
Multi-layer donut plots showing (A) the methodologies used in records analysing Fire-and-IAS (inner ring) and the research aims addressed within each methodology (outer ring) and (B) the methodologies used in records analysing Fire-and-IAS (inner ring), the types of results reported in Fire-and-IAS records (middle ring) and, for the subset of articles with predictive results, the records that used scenario-based approaches (outer ring).

**Figure 7.**
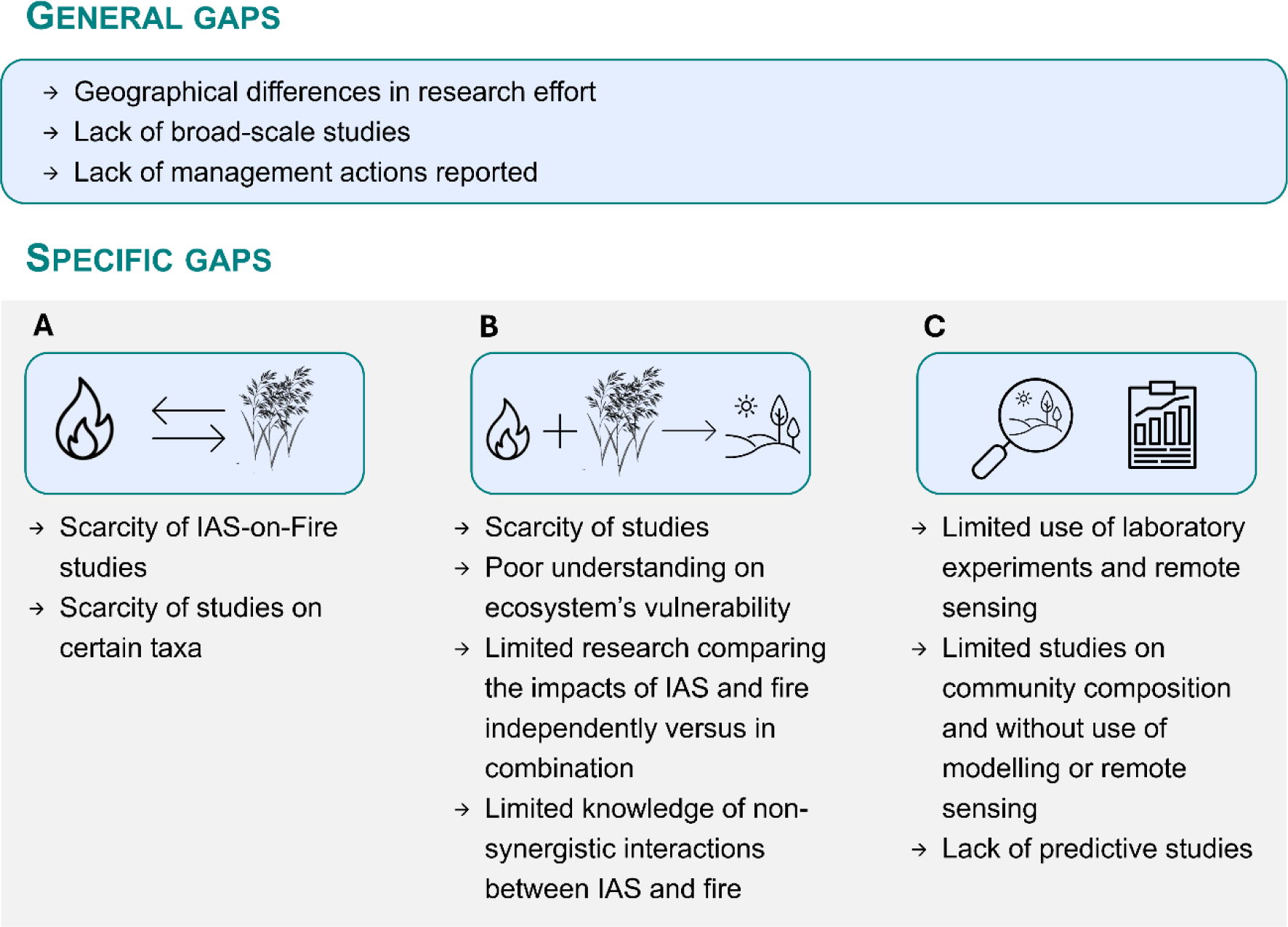
Scheme summarizing knowledge gaps identified in this review. The scheme distinguishes between general gaps (common across all studies) and topic-specific gaps. Topic-specific gaps are grouped into three categories: invasion–fire feedbacks (A), invasion–fire impacts on ecosystems (B), and methods, aims, and result types used in invasion–fire impacts on ecosystem studies (C).

Records about Fire-and-IAS that aim to understand effects on specific species or ecosystem properties are the most prevalent and well represented across all methodological approaches (Figure 6A outer ring). This category includes studies that provide detailed information on individual components of ecosystems, such as reductions in the abundance of species and shifts in ecosystem processes (e.g., McGranahan et al., 2015; Le Maitre et al., 2014), which is the basis for understanding the mechanisms underlying invasion–fire dynamics and linking these findings to broader ecological patterns.

The second most studied aim concerns restoration outcomes (Figure 6A outer ring), either in terms of planning or evaluating them (Kaul et al., 2023; Ricca et al., 2018). These studies employ experimental approaches (Pyke et al., 2022), modelling (Forbis et al., 2006), and remote sensing (Rodhouse et al., 2021) but lack laboratory experiments and purely observational studies. This pattern is consistent with the nature of restoration, which generally requires quantification to guide decision-making and assess effectiveness, typically through ecosystem attributes such as diversity, vegetation structure, and ecological processes (Ruiz-Jaen and Mitchell Aide, 2005). Such studies are particularly important in the current context of increasing priority on biodiversity and ecosystem conservation (UNEP and FAO, 2020; USDA Forest Service, 2018).

Analyses of community composition and IAS management outcomes are the focus of fewer than ten records each (Figure 6A outer ring). Research on IAS management outcomes provides critical information on the effectiveness of control strategies, essential to design conservation plans that must manage multiple threats (Menon et al., 2025). Studies on community composition offer a broader perspective than studies focusing on specific species or ecosystem properties, as community attributes are better suited to inform habitat-specific or system-wide restoration (Timpane-Padgham et al., 2017). Community composition is investigated through experimental studies, observational studies, and laboratory experiments, while IAS management outcomes are examined using experimental studies, observational studies, and modelling.

Regarding result types, most studies focus on explaining invasion-fire interaction (71%; Figure 6B – middle ring). Studies describing the interaction compose 15% of the dataset, although very few are purely descriptive. Most studies with descriptive sections also present explanatory results. This complement is important because, although descriptive data can be important to shape future research by bringing new insights or providing data for other studies, research is more informative and robust when it is grounded on explanatory statistical analysis (King, 2014). Descriptive and explanatory results appear in studies across all methodologies and always occur in similar proportions.

Predictive records account for only 13% of the total and are exclusively obtained through modelling approaches (Figure 6B – middle ring). These records can provide crucial information for management. For example, several records model the impact of IAS on the fuel characteristics of an area and predict the consequences for fire behaviour (Gerber et al., 2024; Tortorelli et al., 2023; McGranahan et al., 2013). Within the subset of predictive records, scenarios are used in 75% (Figure 6B – outer ring). Scenario-based approaches are useful to anticipate how invasion–fire interactions might evolve under future conditions (e.g., climate change) and under different management strategies (e.g., Thomson, 2024; Forbis et al., 2006). This allows stakeholders to understand the potential outcomes of each decision and make informed choices (Peterson et al., 2003). Models and scenarios are essential for understanding, predicting, and managing IAS, guiding prevention and early detection by projecting current and future invasion patterns (Roy et al., 2024). However, models and scenarios of IAS dynamics remain limited, restricting reliable forecasts of future invasions (Seebens et al., 2023).

## 7. Overview, knowledge gaps, and future steps

Several knowledge gaps were found in this review. Some gaps were evident across all topics and related to three main issues: geographical differences in publication, the spatial scale of studies, and biases in the reporting of results.

The research effort is, as expected, concentrated in regions where invasion–fire interactions have pronounced ecological consequences, such as the United States of America and Australia. However, other fire-prone regions remain understudied despite the existence of clear invasion–fire dynamics (Appendix 4). For example, in southern Europe, fire-prone landscapes are often invaded by alien species (Marfella et al., 2023; Morais et al., 2021; Grigulis et al., 2005), yet relatively few studies have explored this topic. Similar patterns are found in Asia where, for example, fire-prone savannas in southeast Asia are also underrepresented in invasion–fire research, despite the documented impacts of inadequate fire regimes and IAS (Tornorsam et al., 2024). In Africa, all research is conducted in South Africa except for two studies in Serengeti, Tanzania. One study concludes that the abundance of IAS was higher in areas of infrequent fire and no fires over a decade (Bukombe et al., 2018) and the other that intense fires can be used to control invasion by *Gutenbergia cordifolia* Benth. ex Oliv. (Mero et al., 2023). Therefore, the lack of studies may partly reflect the resistance of these ecosystems to invasion, as suggested by Visser et al. (2016), but more work is necessary.

Regarding spatial scale, most research has been conducted at local and regional scales or within protected areas, providing valuable detailed insights but often overlooking broader ecological patterns (Appendix 4). Very few studies have addressed invasion–fire interactions at island, national, or international scales. Large-scale approaches, using remote sensing or correlative modelling, can complement local studies by informing, for example, how the realized niche of a native species is affected by IAS and fire (Geyle et al., 2025) or how IAS are distributed according to different fire frequencies (Visser et al., 2016).

Finally, despite the recent attention to management highlighted in the ‘Temporal trends’ section, management actions remain rarely reported. Only a little over 30% of the dataset included any type of management—whether directed at IAS, fire, or broader landscape objectives. Specifically, in IAS-on-Fire studies only 7 address management actions. This lack of management actions reported in the invasion–fire literature could reflect two possibilities. First, few management actions may be implemented in areas affected by invasion–fire interactions. If that is the case, considering the many strong interactions and negative impacts on a wide range of habitats and taxa identified in this review, addressing this gap at both management and research levels should become a priority. Second, management outcomes may be undertaken but remain unpublished due to lack of success. The bias toward publishing positive results is well-documented in several disciplines that inform conservation practice and policy (Wood, 2020). Indeed, unsuccessful management is rarely reported, even though documenting such outcomes is critical for understanding why strategies fail, creating a more balanced evidence base, and improving adaptive management (Catalano et al., 2019). To address this gap, journals should be more receptive to publishing studies reporting “negative evidence”. In addition, creating integrated databases that compile management outcomes and improving the inclusion of grey literature in evidence syntheses could further enhance the availability and diversity of information.

### 7.1. Knowledge gaps per topic

#### Invasion-fire feedback

Among the three directions of interaction, the effects of IAS on fire regimes are the least studied. The scarcity of studies in this direction results in an incomplete understanding of which IAS and which of their traits (e.g., productivity, flammability, phenology) can alter fire regimes. It also hampers our understanding of which components of the fire regime are most affected. This is particularly concerning because many species known to be problematic are underreported or entirely absent from these studies (Appendix 5). This is a critical gap, as understanding these effects is essential to determine whether an invasion–fire regime cycle is established (Brooks, 2008). Invasion-fire cycles, where IAS alter the fire regime and benefit from those changes, can drive regime shifts and generate novel ecosystems in which native species are unable to persist (Fusco et al., 2019; Gaertner et al., 2014).

Plants are the most studied taxa in the context of invasion–fire interactions, likely because they have a direct influence on fire dynamics. In contrast, research on invasive animals is scarce (Figure 3A). Among the few studies examining the effects of fire on animals, 31% report that fire can promote their success (e.g., Atchison et al., 2018; Hradsky et al., 2017; Malumbres-Olarte et al., 2014). Additionally, the only study analysing the potential effects of an invasive animal on fire found that *Sus scrofa* L. reduced fire spread through rooting, which created fire breaks by exposing bare ground and reducing fine fuel loads (Brown et al., 2024). Fungi have been largely overlooked, with only one study addressing IAS-on-Fire interactions where the authors found that the pathogen responsible for Sudden Oak Death increased surface fuel loads in Douglas-fir–tanoak forests in California (Valachovic et al., 2011). These few studies suggest that invasive animals and fungal pathogens can significantly interact with fire and should be further studied.

#### Invasion-fire combined impacts on ecosystems

Research addressing the combined impacts of fire and IAS on ecosystems is limited, comprising only 20% of the dataset. Given that this is a relatively young field, early efforts focused on disentangling how each driver influences the other. However, beyond expanding the knowledge of these interactions, there is now a need to prioritize a more comprehensive and integrative assessment of the ecological consequences.

Regarding ecosystems studied, research has focused on the impacts caused in vulnerable ecosystems, while resilient ecosystems remain largely unexamined. This focus is unsurprising given the need to prioritize conservation efforts under limited resources (Cullen, 2013). However, investigating resilient ecosystems could provide valuable insights into the traits that confer resilience to invasion–fire interactions (Enright et al., 2014), thereby informing strategies for conserving or restoring more vulnerable areas. Comparative studies that assess both vulnerable and resilient ecosystems could be particularly valuable.

The number of studies explicitly comparing the independent and combined impacts of fire and IAS on ecosystems is limited, despite being crucial for understanding their interactive dynamics. For instance, Steidl and Litt (2009) showed that fire effects on small mammal abundance varied with the level of invasion by *Eragrostis lehmanniana*, with a species declining more in burnt invaded areas than in burnt only areas, other increasing in burnt invaded areas and decreasing in burnt only areas, and other with small changes.

Also related to the outcomes of the interaction, publication bias may be limiting our knowledge of non-synergistic invasion-fire interactions. In invasion biology, a context bias exists due to the use of value-laden language, which frames alien species as inherently harmful and predisposes research and management toward emphasizing negative outcomes (Warren et al., 2017). Consequently, invasion–fire studies may disproportionately report cases where IAS and fire exacerbate each other’s effects, while instances in which these disturbances do not reinforce one another, or fire mitigates IAS effects, may be underrepresented. Studies reporting non-synergistic invasion-fire interaction are rare but exist. A study in California’s Big Sur region concluded that wildfire reduced tree mortality caused by the invasive *Phytophthora ramorum*, with higher burn severity linked to lower post-fire disease mortality (He et al., 2021).

#### Methods, aims, and result types in Fire-and-IAS studies

Regarding methodologies used, there is a limited application of laboratory experiments and remote sensing approaches. As previously discussed, studies employing remote sensing demonstrate its potential to capture invasion–fire dynamics across large spatial and temporal scales (e.g., Rees et al., 2024; Visser et al., 2016; Setterfield et al., 2013), making it a valuable tool to address the previously noted gap of broad-scale and long-term observational studies. Laboratory experiments, on the other hand, provide the advantage of controlling variables within this complex interaction. Fire-related factors such as smoke or intensity fluctuations can be manipulated to test their effects on native species and IAS (e.g., Figueroa and Cavieres, 2012; Clarke and French, 2005). Such experiments can identify physiological thresholds and be used to design management strategies that leverage the strengths of native species while targeting the vulnerabilities of invaders. Moreover, both methodologies rely on standardized procedures, which facilitates comparisons among case studies and ensures that differences in outcomes reflect the characteristics of the studied species or ecosystems.

Research on community-level responses to invasion–fire interactions remain scarce. This is a major gap since community-level dynamics is one of the most used approaches to assess the resilience of ecosystems to disturbances (Capdevila et al., 2021). Moreover, from the set of studies focused on community composition, none use modelling or remote sensing methodologies (Figure 6B). This limits opportunities to generalize findings across systems or to scale them up to the landscape level.

Finally, the number of predictive approaches is limited, which constrains our ability to anticipate how invasion–fire dynamics may unfold under different scenarios (e.g., climate change, land-use change, management decisions). Moreover, the predictive studies that do exist are often restricted to specific species or regions, which hinders generalizing their conclusions. Expanding predictive modelling efforts would provide decision-makers with more robust tools to prepare for and mitigate future risks.

In sum, future research should prioritize informing management, increasing understanding of IAS-on-fire, and improving knowledge of case-specific invasion–fire interactions and their outcomes. It is also necessary to focus on underrepresented regions (e.g., Europe, Africa), broaden taxonomic coverage (e.g., animals, fungi), expand the spatial scale of studies, and promote multi-scale integrative research addressing interactions across drivers and scales. To achieve this, it will be crucial for future studies to diversify their objectives and methodologies.

## 8. Conclusions

- The present review is the first to assess invasion-fire interactions across all directions of interaction, all taxonomic groups, and with a focus on their combined ecosystem impacts.
- Although the field is relatively recent, several key patterns are already well established: fire generally promotes IAS, IAS can modify fire regimes and lead to regime shifts, and their combined impacts on ecosystems are mostly negative.
- Several knowledge gaps were found. Invasion-fire interactions are complex and not always synergistic, highlighting the need for more nuanced approaches to study their impact on ecosystems. Research also remains underrepresented for some taxa and regions and with few studies addressing community-level responses or broad-scale and multi-scale dynamics. The outcomes of management actions are rarely reported, but when implemented through prescribed fire, positive impacts on IAS are most often observed.
- Future studies should (1) increase the understanding of IAS effects on fire, (2) improve knowledge of case-specific invasion–fire interactions and their ecosystem effects, and (3) test and refine management actions. Strengthening the link between research and management is essential to develop strategies that mitigate the negative impacts of invasion–fire interactions and enhance ecosystem resilience.

## Acknowledgements

CG Lima is supported by the FCT - Portuguese Foundation for Science and Technology through the 2021 PhD Research Studentships [grant reference 2021.05661.BD]. A Regos is currently funded by the ‘Ramon y Cajal’ fellowship program of the Spanish Ministry of Science and Innovation (RYC2022-036822-I).

## Notes

### Competing Interest Statement

The authors have declared no competing interest.

