## Appendix 1 for "Combined impacts of invasive alien species and fire on ecosystems are complex, mostly negative, and understudied: a global review"

Table 1 - Keyword string used to perform the literature search and references used for their selection.

| Filter | IAS | Fire regime | Interaction |
| --- | --- | --- | --- |
| Keywords | "ecological invasion*" OR "biological invasion*" OR "invasion* biology" OR "invasion* ecology" OR "invasive species" OR "alien species" OR "nonnative species" OR "non-native species" OR "nonindigenous species" OR "non-indigenous species" OR "allochthonous species" OR "exotic species" OR "invader*" OR "introduced species" OR "invasive*" OR "introduced landscape" OR "non-native landscape" OR "nonnative landscape" OR "nonindigenous landscape" OR "non-indigenous landscape" OR "allochthonous landscape" OR "novel ecosystem" | ("fire regime*" OR "fire intensity" OR "fire size" OR "fire extent" OR "fire seasonality" OR "burn probability" OR "fire return interval" OR "fire severity" OR "fire frequency" OR "fire management" OR "fire occurrence" OR "fire event*" OR "fire ecology" OR "fire history" OR "fire interval" OR "burning treatments" OR "fire behavio*r" OR "fire type" OR "fire rotation") | ("interact*" OR "impact*" OR "*ffect*" OR "fav*r*" OR "promot*" OR "benefit*" OR "feedback" OR "influence*" OR "alter*" OR "relation*" OR "persist*" OR "suppress*" OR "facilitate*" OR "respon*" OR "dynamic*" OR "co-occur*" OR "modif*" OR "deviat*" OR "consequence*" OR "disturb*" OR "displacement" OR "implicat*" OR "damage*" OR "disrupt*" OR "chang*") |
| References | Essl et al. (2018); Roy et al. (2024) | Kelly et al. (2023); Krebs et al. (2010) | Brooks et al. (2004; Tomat-Kelly and Flory (2023) |
